# Tissue-Specific Cell Type Annotation with Supervised Representation Learning using Split Vector Quantization and Its Comparisons with Single-cell Foundation Models

**DOI:** 10.1101/2024.12.09.627458

**Authors:** Yusri Dwi Heryanto, Yao-zhong Zhang, Seiya Imoto

## Abstract

Cell-type annotation in single-cell data involves identifying and labeling the cell types based on their gene expression profiles or molecular features. Recently, with advances in single-cell foundation models (FMs), unsupervised annotation and transfer learning with FMs have been explored for cell-type annotation tasks. However, because FMs are usually pre-trained in an unsupervised manner on data spanning a wide variety of tissues and cell types, their representations for specific tissues may lack specificity and become overly generalized. In this work, we propose a novel supervised representation learning method using split-vector-quantization, single-cell Vector-Quantization Classifier (scVQC). We evaluated scVQC against both supervised and unsupervised representation learning approaches, with a focus on foundation models pretrained on large-scale single-cell datasets, such as scBERT and scGPT. The experimental results highlight the importance of label supervision in cell-type annotation tasks and demonstrate that the learned codebook effectively profiles and distinguishes different cell types.

## 1. Introduction

Cell annotation is a crucial step in single-cell analysis. Two commonly used methods for this are unsupervised clustering with marker-based annotation and label transfer using reference datasets (Clarke et al. [2021]). In the first method, cells are grouped into clusters based on their expression profiles using unsupervised clustering algorithms. These clusters are then annotated by identifying differentially expressed genes and matching them to known cell type markers from literature or databases. The unsupervised clustering methods can be time-intensive and requires expertise in selecting reliable marker genes. To overcome these limitations, label transfer methods were developed. This approach utilizes annotated reference datasets to automatically assign cell type labels to new data through computational tools, offering a faster, consistent solution that leverages prior knowledge from reference datasets.

Recent studies demonstrated the effectiveness of foundation models (FMs) in single-cell analysis (Yang et al. [2022], Cui et al. [2024a], Szalata et al. [2024]). These models are pretrained on extensive single-cell datasets spanning various human tissues, organs, and even data from other species. After pretraining, the models can be fine-tuned for a range of downstream tasks. Among the leading LLMs in single-cell analysis are scGPT (Cui et al. [2024a]) and scBERT (Yang et al. [2022]).

While FMs hold significant promise for single-cell genomics, the field also faces notable challenges and criticisms. Since FMs are pre-trained in a self-supervised or unsupervised manner on diverse single-cell datasets from various tissues, the representations they learn may lack specificity. This issue arises because the definition of a cell type often depends on the tissue or organ in which it resides (e.g., epithelial cells in the skin differ from epithelial cells in the pharynx). As a result, FMs may generalize too broadly, potentially compromising their ability to capture tissue-specific distinctions (Zeng [2022]). Additionally, the computational, time, and memory demands of FMs are considerable, even for fine-tuning only (Naveed et al. [2023], Kedzierska et al. [2023]). Some FM architectures also require specific GPU configurations; for instance, scGPT relies on FlashAttention (Dao et al. [2022]), which is unavailable on older or smaller GPUs.

In cell type annotation, foundation models (FMs) map input features into an embedding space learned from pretrained data, where a classifier is trained to categorize cells. Cell representations in this embedding space is learned in an unsupervised manner. A common approach involves using a variational autoencoder (VAE) to learn cell representations (Gomari et al. [2022]). However, standard VAEs have limitations in single-cell type classification, as they assume a multidimensional normal prior for the low-dimensional latent variables. This assumption often leads to clustering cell representations near the center of the latent space, even when distinct cell types are present (Ding and Regev [2021]). Additionally, VAE models are prone to degenerate solutions, a problem known as posterior collapse (Dai et al. [2019]). To address these issues, the Vector Quantised-Variational AutoEncoder (VQ-VAE) was introduced (Oord et al. [2017]). Unlike VAEs, VQ-VAE encodes discrete rather than continuous codes. Discrete representations are generally more suitable for cell type prediction, providing clear boundaries between cell types, reduced overlap in latent space, and avoiding posterior collapse (Cui et al. [2024b]).

However, representations learned during self-supervised pretraining or unsupervised learning may not be optimal for cell classification. A supervised neural network can be trained to refine these representations for classification tasks. For instance, SCLSC (Heryanto et al. [2024]) uses supervised contrastive learning to enhance cell type representations, aligning each sample closely with its representative cell in the new embedding space while maintaining separation from representatives of other labels. SCLSC assumes that the gene profile of a representative cell for each type is the average of gene profiles for samples within that cell type.

Inspired by the previous works, we developed the single-cell Vector-Quantization Classifier (scVQC) model for annotating cell types in scRNA-seq data. The scVQC model leverages split-vector-quantization to create discrete representations of the cells, which enhances the distinction between cell types by capturing essential features while reducing noise. Split-vector quantization (Kobayashi et al. [2022]) improves codebook utilization by dividing vectors into smaller, equally sized subvectors and quantizing them individually using a shared codebook. Studies show this approach outperforms conventional vector quantization in utilizing the codebook more effectively (Cui et al. [2024b]). Further, we incorporate a supervised clustering approach that leverages prior knowledge of labeled cell types to guide the clustering process. Unlike traditional clustering methods, which rely solely on intrinsic data structures, supervised clustering assumes that data points have predefined class labels and seeks to identify high-density clusters that predominantly consist of a single class (Eick et al. [2004]). By training on annotated datasets, scVQC improves cluster assignment accuracy, ensuring biologically meaningful classifications and reducing ambiguity in cases where closely related cell types exhibit overlapping gene expression profiles.

In this study, we conducted a comprehensive comparison of foundation model (FM)-based methods, specifically scGPT and scBERT, with supervised neural network approaches, including SCLSC and scVQC. Utilizing four benchmark single-cell datasets, we evaluated the performance of these methods across multiple dimensions: classification accuracy, clustering quality, representation learning capabilities, and scalability (Figure 1). Our analysis aimed to provide insights into how well each approach captures the underlying structure of single-cell data, as well as their ability to handle large-scale datasets efficiently.

**Fig. 1:**
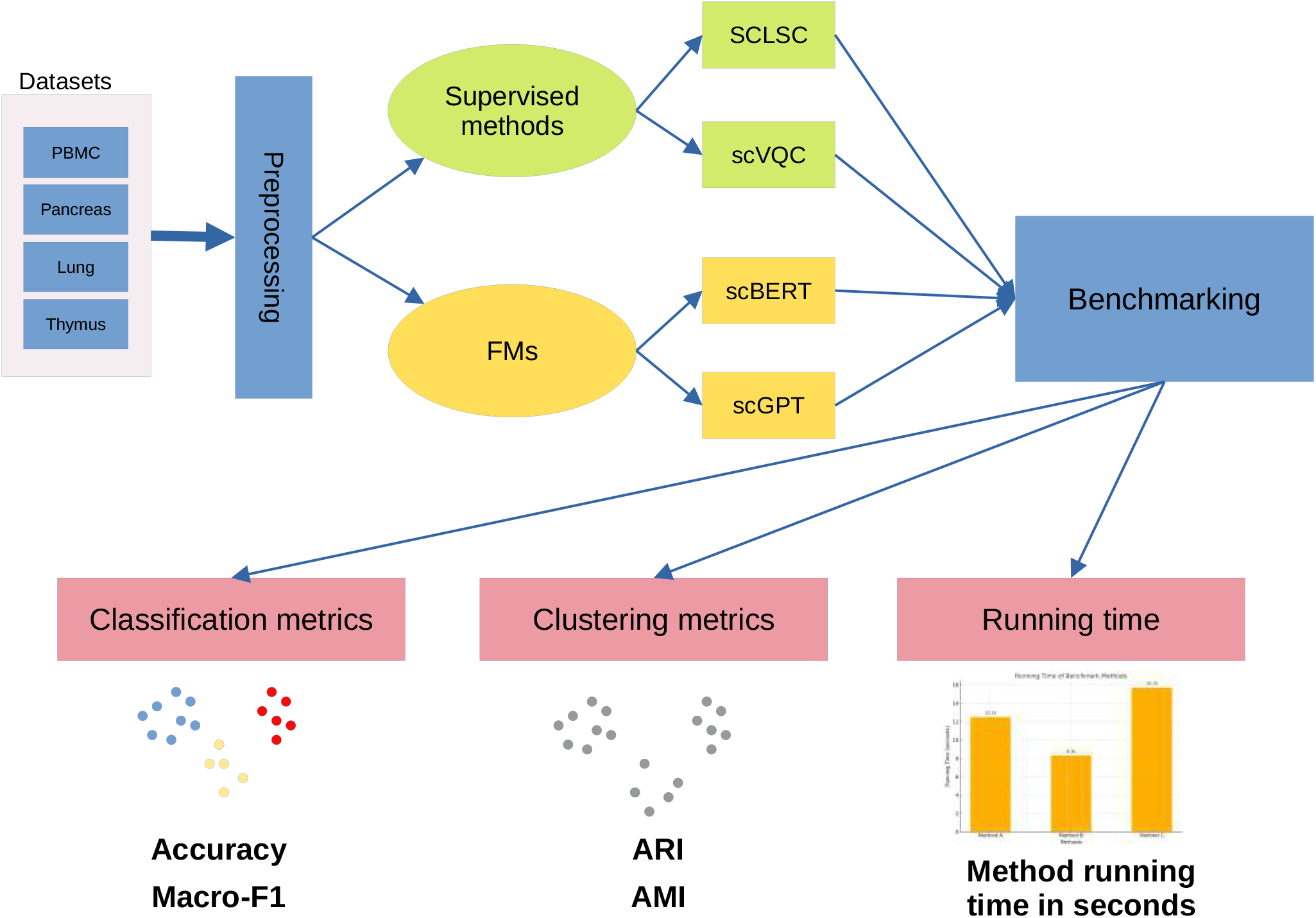
The workflow of our study. The analysis begins with preprocessing four datasets (PBMC, Pancreas, Lung, and Thymus), followed by comparisons between supervised methods (SCLSC and scVQC) and foundation models (scBERT and scGPT). Benchmarking includes evaluations of classification metrics (Accuracy, Macro-F1), clustering metrics (ARI, AMI), and method running times.

This study contributes to the field of single-cell genomics in the following ways:

1. It introduces scVQC, the first model to incorporate split-vector quantization for annotating cell types in single-cell RNA-sequencing (scRNA-seq) data.
2. Through a comparative analysis of foundational models (scGPT, scBERT) and supervised neural network-based methods (SCLSC, scVQC), the study demonstrates that supervised learning approaches achieve performance that is comparable to, or even exceeds, that of foundational models, while offering faster processing and more efficient computational requirements.
3. It introduces the use of the feature spectrum to obtain a novel representation of cell types by applying the TF-IDF transformation to the vector quantization codebook corresponding to each cell type.

## 2. Materials and methods

### 2.1. Data Preprocessing

Before analysis, we split each dataset into two parts: 60% for the training set and 40% for the test set using a stratified split. We then applied a preprocessing pipeline on these datasets. In this pipeline, cells with high mitochondrial gene expression (greater than 5% of the total cell count) were excluded, along with cells showing minimal gene expression (fewer than 200 genes per cell) and genes detected in only a small number of cells (fewer than 3 cells expressing the gene). Afterward, we normalized each cell’s count to 10,000 counts and applied a log(x+1) transformation. We then selected the top 3,000 highly variable genes (HVGs) for further analysis.

### 2.2. The scVQC Framework

The scVQC takes as input a preprocessed gene-profile matrix *X* of dimensions *n* × *d*, where *n* represents the number of cells and *d* denotes the number of genes. The model is composed of three key components: an encoder that maps the feature space to a continuous latent embedding space, a split quantizer that converts this continuous space into a discrete latent embedding space, and a classifier head that predicts cell types based on these discrete embeddings (Figure 2).

**Fig. 2:**
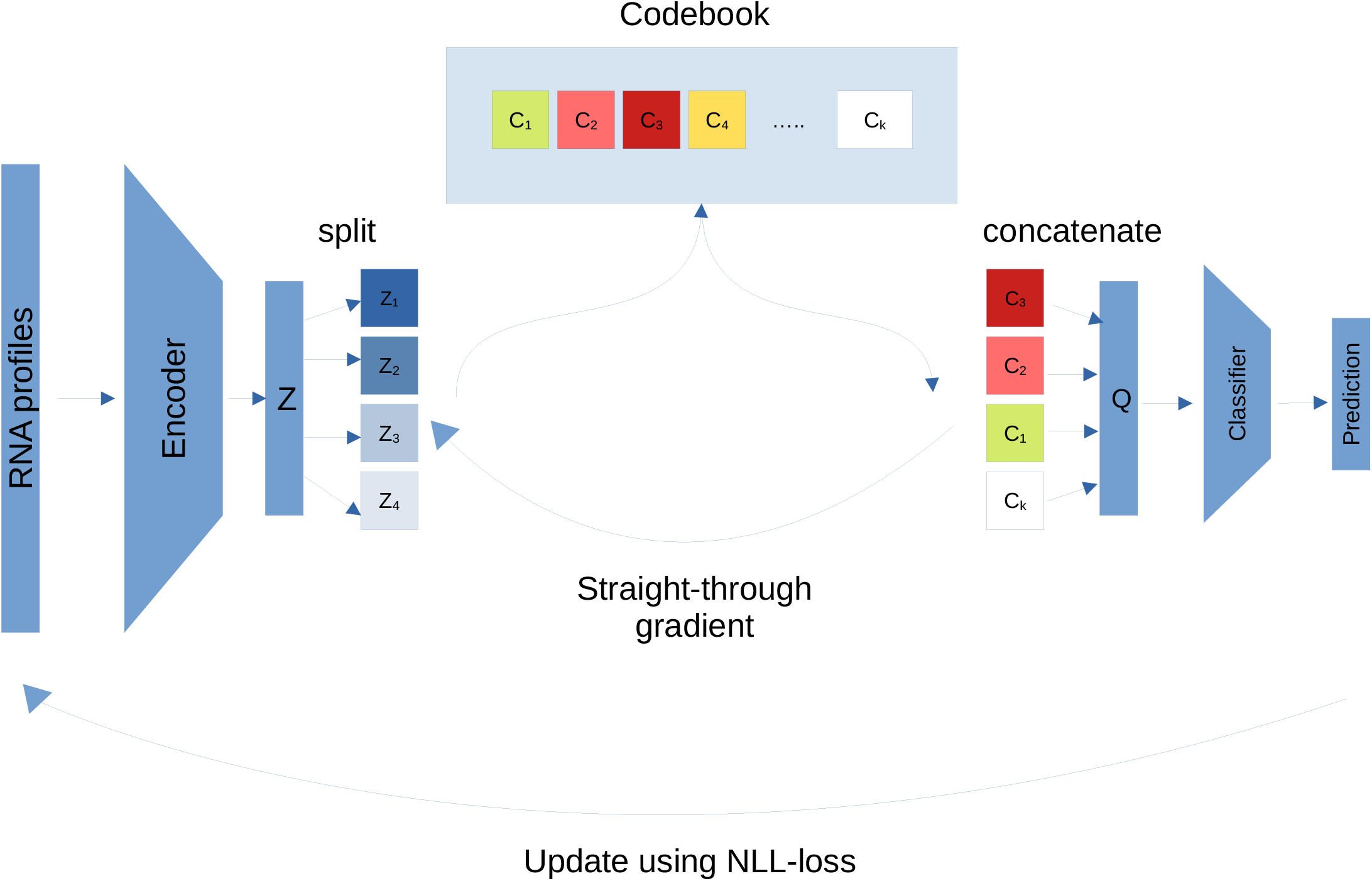
The scVQC architecture.The scVQC consists of three main components: a 3-layer MLP encoder, a split-vector quantizer, and a dense layer with a log-softmax activation serving as the classifier head.

#### 2.2.1. Encoder structure

The encoder accepts *x*_*i*_ ∈ ℝ^*d*^ as the input where *d* denotes the number of genes. The gene profile *x*_*i*_ represents the *i*-th row of the matrix *X*, which is a vector of gene expression values for cell *i*. The encoder in scVQC consists of two Linear-dropout-Relu-batch normalization block layers and followed by dense linear layer. The output of the encoder for each *x*_*i*_ is *z*_*i*_ ∈ ℝ^*d′*^ where *d*^*′*^ denotes the dimension of embedding space.

In our study, the linear layers contain 1024, 256, and 100 nodes in the first, second, and third layers, respectively. The dropout rate for both dropout layers is set to 0.1.

#### 2.2.2. Split quantizer

The latent embedding *z*_*i*_ is then split into *M* vectors 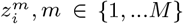. Using Vector Quantization (Oord et al. [2017]), each split 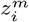 is assigned to the nearest code *e*_*k*_ ∈ R^*d*\*^ , *k* ∈ {1, …*K*} in the codebook. This assignment is represented by as 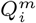.

Here 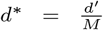 is the dimension of the codes and K is a hyperparameter defining the number of codes within the codebook. This assignment is formulated as follows:

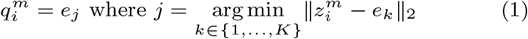

Next, we concatenate back each 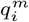 into *q*_*i*_ ∈ R^*d′*^ that represents the quantized *z*_*i*_. The

The split vector quantization (Kobayashi et al. [2022]) is a modified approach to vector quantization designed to improve codebook utilization by addressing the issue of underused codes. Instead of quantizing all dimensions of a vector at once, the vector is first divided into smaller, equally sized subvectors. Each subvector is then quantized individually using a shared codebook. From previous studies, this method enhanced codebook utilization compared to models that perform quantization on the entire vector simultaneously (Cui et al. [2024b]).

In our study, we set the codebook to have 200 codes, an embedding dimension of 100, and 20 splits (i.e. resulting in each split having a dimensionality of 100 ÷ 20 = 5).

#### 2.2.3. Classifier head

The classifier head accepts *q*_*i*_, the output from the split quantizer, as its input. It consists of a linear layer followed by a log-softmax activation layer for classification.

### 2.3. Training Process

For each cell, a gene expression profile vector *x*_*i*_ and its associated cell type label *y*_*i*_ from the training dataset are provided as input to scVQC. The encoder transforms the input data into a continuous latent embedding, which is then converted into a discrete latent embedding by the split quantizer. This discrete latent embedding is subsequently used by the classifier head to predict the cell type. The training objective of scVQC comprises three components. The first component is the negative log likelihood (NLL) loss for measuring classification accuracy, defined as:

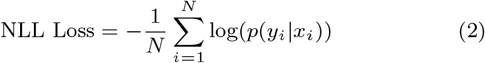

where *N* is the number of samples, and *p*(*y*_*i*_|*x*_*i*_) is the predicted probability for the correct class *y*_*i*_, given the input*x*_*i*_. This classification loss updates the encoder and classification head.

The second component is the commitment loss. This loss ensures the encoder output is “committed” to a specific code in the codebook. The commitment loss defined as follows:

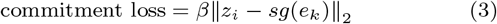

where *sg* represents the stop-gradient operator, which acts as an identity function during forward computation but has zero partial derivatives. This operator prevent the operands get updated during backpropagation. The commitment loss only update the encoder. The *β* is a factor that balances the trade-off between the commitment loss and other losses. We set *β* = 0.25, the same default value used in the original paper (Oord et al. [2017]).

The last component is the codebook update, designed to update the codebook by moving selected code vectors closer to the encoder’s output. We used the Exponential Moving Average (EMA) from Oord et al. (Oord et al. [2017]) as the function for updating the codebook. Specifically, for each code vector *e*_*k*_ in the codebook, the EMA update rule is:

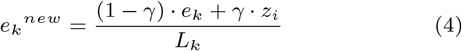

where *γ* is the decay factor of EMA set at 0.99, *e*_*k*_^*new*^ and *e*_*k*_ is the updated and old codes, *z*_*i*_ the encoder’s output for an input that has been assigned to *e*_*k*_, and *L*_*k*_ is the usage count for *e*_*k*_. This count helps normalize updates and prevent inactive codebook vectors from drifting randomly.

Thus, the total training loss:

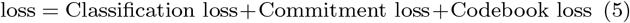

We trained the scVQC for 5 epochs in our study with learning rate 5e-4 using Adam optimizer. For datasets containing more than 100,000 cells, the batch size is set to 128. For datasets with 100,000 cells or fewer, the batch size is set to 32.

### 2.4. Competing Methods

#### 2.4.1. SCLSC

SCLSC (Heryanto et al. [2024]) is a deep learning method developed to apply supervised contrastive learning principles for transferring annotations from a single-cell reference dataset to a query dataset. By contrasting each cell with its cell type representation, SCLSC effectively learns a low-dimensional representation that is well-suited for cell type prediction. We used the default hyperparameters provided in the SCLSC GitHub repository (https://github.com/yaozhong/SCLSC), with a maximum training epoch of 30.

#### 2.4.2. scGPT

scGPT (Cui et al. [2024a]) is a generative pre-trained model designed for analyzing single-cell RNA data. Its core architecture consists of stacked transformer layers with multi-head attention, enabling simultaneous generation of cell and gene embeddings. Pretrained on large-scale, diverse datasets with over 33 million cells from the CELLxGENE collection, scGPT produces rich cell representations. For cell type annotation, scGPT offers two methods: zero-shot and fine-tuning. For the zero-shot approach, we followed the hyperparameters and workflow outlined in the scGPT reference mapping tutorial available on GitHub (https://github.com/bowang-lab/scGPT/blob/main/tutorials/Tutorial_Reference_Mapping.ipynb). For the fine-tuning approach, we adhered to the instructions provided in the scGPT annotation tutorial (https://github.com/bowang-lab/scGPT/blob/main/tutorials/Tutorial_Annotation.ipynb).

#### 2.4.3. scBERT

scBERT (Yang et al. [2022]) is a large language model built on the architecture and pretraining strategies of BERT (Bidirectional Encoder Representations from Transformers) (Devlin et al. [2018]). It was pretrained on the PanglaoDB dataset, which includes 1,126,580 cells from 74 different tissues, gathered from a range of experimental sources and platforms. After pretraining, scBERT is fine-tuned specifically for cell-type classification. The query dataset annotation was carried out using the default hyperparameters and workflow provided in the scBERT repository (https://github.com/TencentAILabHealthcare/scBERT). To obtain the cell embeddings, we used the output generated by the PerformerLM module of scBERT.

### 2.5. Profiling Cell Types Using Feature Spectrum

The feature spectrum was obtained using the Term Frequency Inverse Document Frequency (TF-IDF) transformation applied to the utilization frequencies of codes within the codebooks. In this approach, each code in the codebook was treated as a word, each cell type as a document, and the entire single-cell dataset as a corpus or collection of documents. To calculate the Term Frequency (TF), we measured the relative frequency of each code within a cell type by dividing the raw count of a code’s occurrences in the cell type by the total number of codes in that cell type. To evaluate how common or rare a code was across all cell types, we computed the Inverse Document Frequency (IDF) as the logarithm of the inverse of the fraction of cell types (documents) containing the code. The TF-IDF for each code in each cell type was then calculated as *TF* × *IDF* . This process was applied to all codes across all cell types. It will produce a TF-IDF matrix *T* ∈ *R*^*C*×*Q*^ where *C* is the number of cell types and *Q* the number of codes. Each row of the matrix *T* is the feature spectrum of the corresponding cell type.

To aid the visualization of the matrix *T* , we rearranged its columns as follows. First, for each column *j* we computed two measures:

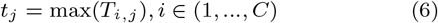

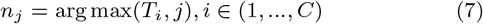

where *C* is the number of cell types. The columns of the *T* were then reordered in non-descending order based on the *n*_*j*_ values, grouping codes that are highly utilized within each cell type. The last step, for each row, we selected the columns that contained *t*_*j*_ . The selected columns were then further reordered in non-ascending order of the *t*_*j*_ values. This final step was repeated for each row.

### 2.6. Unseen cell experiment

We conducted stratified random splits on the PBMC datasets, dividing them into training, validation, and test datasets in an 6:4 ratio. CD19+ B cells were then excluded from the training and validation datasets. The training dataset was used as a reference to predict the annotations of the test dataset using scVQC.

### 2.7. Evaluation Metrics

We assessed the performance of our method relative to other approaches using Accuracy and Macro-Weighted F1 Score metrics. To compute these scores, we calculated the True Positives (TP), True Negatives (TN), False Positives (FP), and False Negatives (FN). Accuracy was defined as the ratio of correct classifications, both positive and negative, to the total number of classifications. The accuracy was calculated as follows:

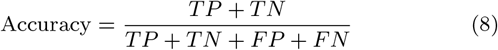

For the macro-weighted F1-score, we calculated precision *P*_*i*_ and recall *R*_*i*_ for each cell type *i* ∈ *C* to obtain the F1 score for each cell type. We then computed the unweighted average of these F1 scores. The formula is as follows:

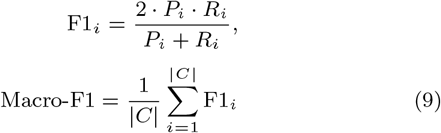

where |*C*| is the number of cell types. We used the sklearn implementations sklearn.metrics.accuracy score() and sklearn.metrics.f1 score(average=‘macro’) for calculating accuracy and macro-F1 score, respectively.

We further evaluated our method’s ability to learn low-dimensional representations using clustering metrics, including the Adjusted Rand Index (ARI) and Adjusted Mutual Information (AMI). Both ARI and AMI are adjusted versions of the Rand Index and Mutual Information scores, accounting for chance and correcting for agreements that may occur solely by chance between clusterings. To compute these metrics, we used the implementations provided by sklearn.metrics: adjusted rand score() for ARI and adjusted mutual info score() for AMI.

## 3. Results

### 3.1. Classification performance comparison

We conducted a comparative analysis of cell type prediction performance across four human datasets: PBMC 68k (Zheng et al. [2017]), lung (Luecken et al. [2021]), thymus (Park et al. [2020]), and pancreas (Luecken et al. [2021]). We compared scVQC with a supervised model and two foundational models. The supervised model, SCLSC (Heryanto et al. [2024]), uses supervised contrastive learning to learn cell representations in a low-dimensional embedding space optimized for classification. The foundational models are scBERT, a BERT-based large language model (Devlin et al. [2018]) pre-trained on 1 million human cells for cell type prediction, and scGPT (Cui et al. [2024a]), a foundation model for single-cell biology pre-trained on over 33 million single-cell sequencing data, supporting diverse downstream tasks such as cell-type annotation, multi-omic integration, and gene network inference.

The supervised approaches (scVQC and SCLSC), particularly scVQC, consistently outperform the foundational models in key metrics across most datasets (Figure 3). For accuracy, scVQC achieves the highest scores in the lung, pancreas, and thymus datasets, while SCLSC also performs competitively, maintaining scores close to scVQC. Among the foundational models, scGPT-zero-shot performs poorly in accuracy, highlighting its limitations without dataset-specific training. In contrast, scGPT-fine-tuned shows significant improvement, with accuracy reaching 0.8280 for PBMC and 0.9460 for pancreas, though it still lags behind supervised methods. scBERT performs well in accuracy, achieving the highest score for PBMC (0.860) and competitive results in other datasets, though it generally falls short of scVQC’s consistency.

**Fig. 3:**
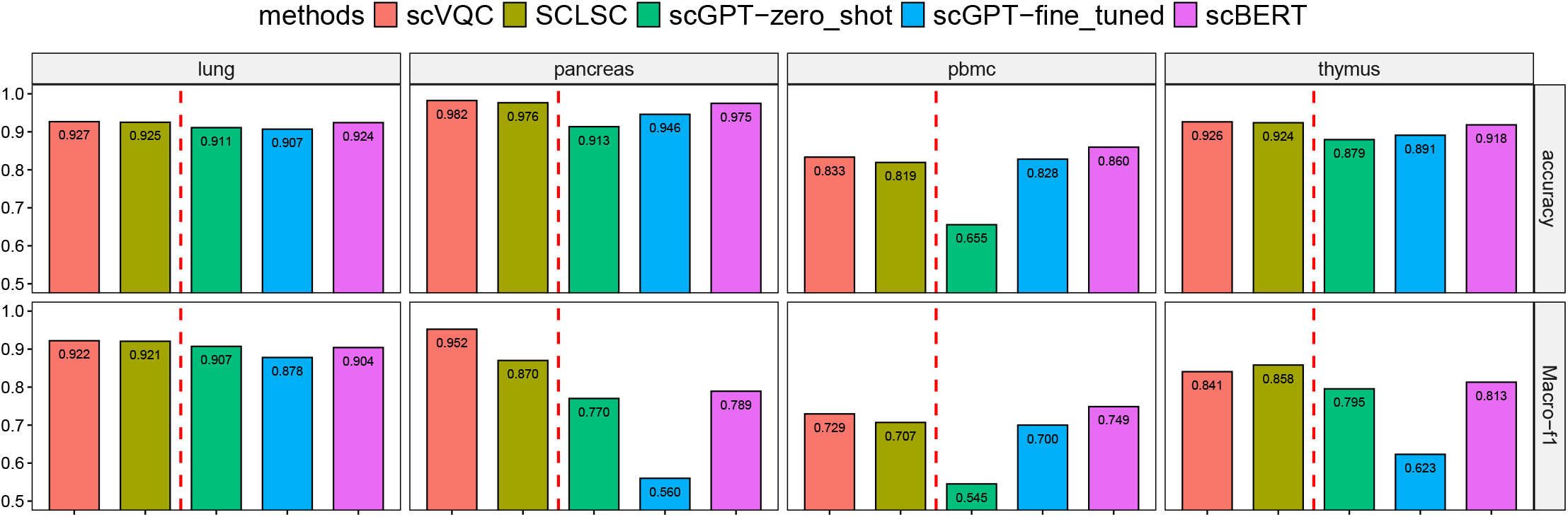
Comparison of Cell Classification Performance. The supervised approaches (scVQC and SCLSC; left of the red dashed line) have better classification performance overall compared to foundational models (scBERT and scGPT; right of the red dashed line) for cell classification tasks using accuracy and Macro-F1 metrics.

For the Macro-F1 metric, which evaluates the balance between precision and recall, the supervised approaches again show clear advantages (Figure 3). scVQC excels with the highest Macro-F1 scores in the pancreas and thymus datasets, demonstrating its ability to classify both common and rare cell types effectively. SCLSC follows closely in performance but lags behind scVQC, particularly in the pancreas dataset. In contrast, the foundational models demonstrate weaker performance. scGPT-zero-shot struggles the most, with notably lower Macro-F1 scores such as 0.5452 for PBMC and 0.7700 for pancreas, indicating its inability to balance precision and recall without dataset-specific fine-tuning. While scGPT-fine-tuned shows improvements, it still falls short in datasets like pancreas, highlighting its limitations even after fine-tuning. scBERT, though competitive in PBMC and lung, is generally outperformed by the supervised approaches, particularly in challenging datasets like pancreas and thymus. These results highlight the superior performance of supervised methods in learning task-specific representations and effectively handling imbalanced cell types, compared to the foundational models.

Overall, supervised approaches, especially scVQC, exhibit superior performance in both accuracy and Macro-F1 compared to foundational models. scVQC’s ability to learn task-specific cell representations through supervision gives it a distinct edge over foundational models, which, despite their generalizability, underperform in some tasks.

### 3.2. Evaluation of clustering metrics

We further examined scVQC and other methods ability to cluster cell representations in the lower-dimensional embedding space. We used ARI and AMI metrics to assess clustering quality, with higher scores indicating a closer alignment with true cell type labels.

Our results reveals a significant advantage for supervised methods (Figure 4a). scVQC consistently demonstrates superior performance, achieving the highest ARI scores in PBMC, lung, and pancreas, while also excelling in AMI with top scores in lung and pancreas. Although SCLSC performs slightly better than scVQC in thymus for both ARI and AMI, it generally remains second to scVQC. In contrast, scGPT-zero-shot struggles with poor clustering performance, reflected in low ARI and AMI scores across all datasets. scGPT-fine-tuned shows significant improvement, particularly in lung, but it still falls short of the supervised approaches, especially in pancreas and pbmc. scBERT shows the weakest performance overall, with consistently low scores across nearly all datasets, highlighting its significant limitations in clustering tasks. Overall, supervised approaches, particularly scVQC, are far more effective in clustering cells and aligning with true cell types compared to foundational models, even when the latter are fine-tuned (Figure 4b).

**Fig. 4:**
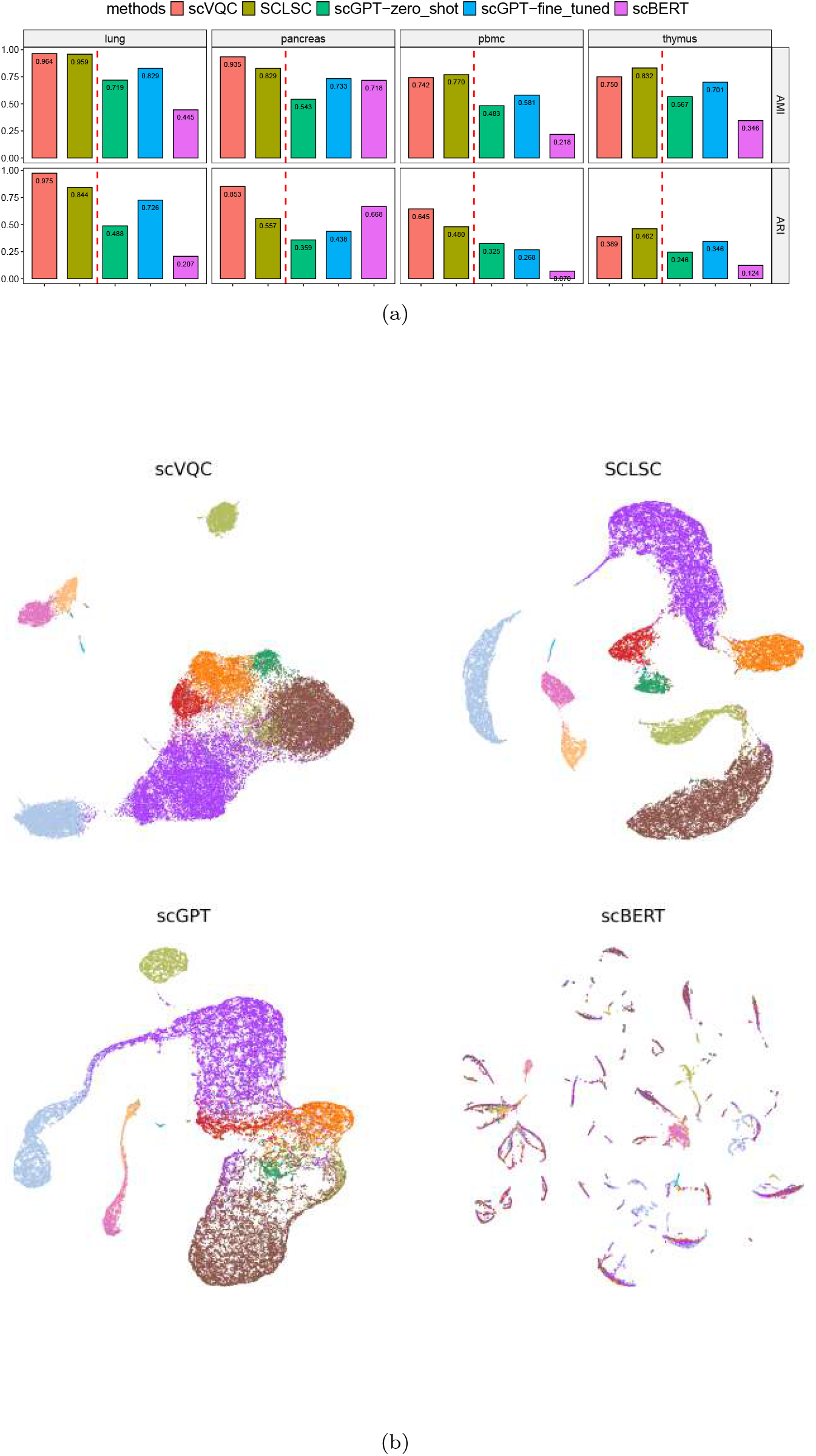
**(a)**Supervised approaches (scVQC and SCLSC; left of the red dashed line), consistently outperform FMs (scGPT and scBERT; right of the red dashed line) on every dataset, as reflected by higher Adjusted Rand Index (ARI) and Adjusted Mutual Information (AMI) scores. **(b)** The UMAP visualization of PBMC cells in the embedding space learned by the benchmarking methods, with colors representing different cell types. The supervised approaches, scVQC and SCLSC, generate embeddings where cells of the same type are more tightly clustered and distinctly separated from other cell types, compared to the foundational models, scGPT and scBERT.

### 3.3. scVQC can learn the representation of the unseen cell in PBMC dataset

scVQC is designed to transfer label annotations from a reference dataset to a query dataset. However, this process becomes more challenging when the query dataset contains cell types that are absent from the reference dataset, limiting scVQC’s ability to predict their annotations accurately. Nevertheless, scVQC is capable of learning meaningful representations for these unseen cell types. To evaluate its performance in such scenarios, we conducted an experiment using the PBMC dataset. Specifically, CD19+ B cells were excluded from the training dataset. Using this training dataset, scVQC learned the representations of each cell in the test dataset. Remarkably, scVQC was able to project the unseen CD19+ B cells as distinct from other cell types (Figure 5). This highlights scVQC’s potential to infer representations of unseen cell types from incomplete data.

**Fig. 5:**
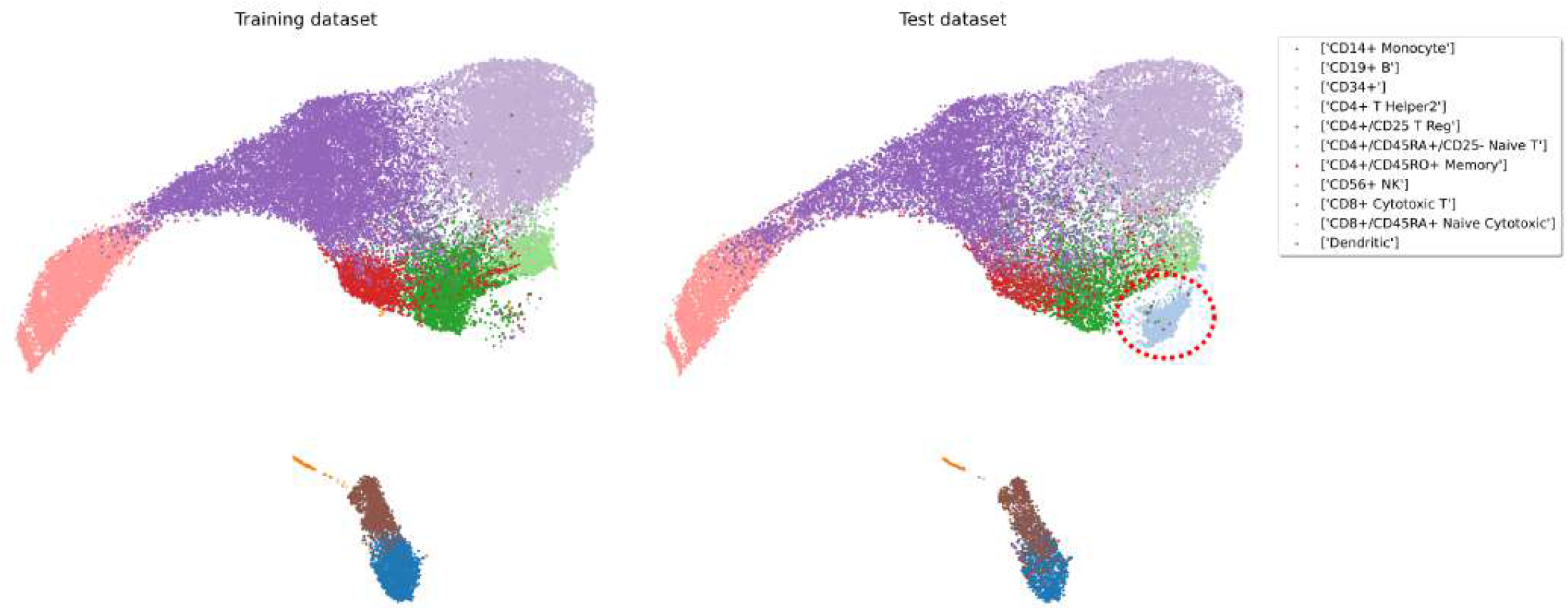
Representation of Unseen Cells. The scVQC demonstrates its ability to infer representations of unseen cells from an incomplete training dataset (left). In the test dataset (right), the model effectively identifies and projects previously unseen CD19+ B-cells (highlighted within the red circle), maintaining a clear separation from other cell types in this UMAP projection.

### 3.4. Evaluation of running time

In this experiment, we measured the running time of scVQC and other methods on an NVIDIA V100 GPU. We used the thymus dataset, which contains over 200,000 cells, making it suitable for benchmarking. Running time is defined as the total time required for training the model and predicting cell types. In our study, scVQC is the fastest, completing the task in just 48.06 seconds. SCLSC follows, taking 377.51 seconds, making it significantly slower than scVQC but faster than the other methods. Among the foundational models, scGPT-zero-shot has a reasonable runtime of 297.35 seconds, reflecting its lighter computational requirements compared to fine-tuned models. However, scGPT-fine-tuned exhibits a substantial increase in runtime, taking 4768.43 seconds, which highlights the additional computational cost associated with fine-tuning. scBERT is by far the slowest, with an extensive runtime of 70493.27 seconds, underscoring its inefficiency and impracticality for applications requiring rapid processing. Overall, scVQC stands out as the most efficient method, while fine-tuned and large foundational models like scGPT and scBERT are significantly slower, limiting their practicality in time-sensitive tasks.

### 3.5. Feature spectrum

The feature spectrum is a novel representation of cell types, created by applying the TF-IDF transformation to the utilization of codes within each cell type. Drawing an analogy to natural language processing, the codes in the codebooks are analogous to words, cell types correspond to documents, and the dataset represents a corpus or collection of documents. In this work, we applied the feature spectrum to identify characteristics of cell types in the PBMC dataset.

Figure 6 illustrates a distinct set of specific features that are either enriched or depleted within the feature spectrum of each cell type in the pancreas dataset. The cell-type-specific feature spectrum, derived from discrete cell embeddings of the single-cell RNAseq profiles, offers an interpretable method for visualizing the overarching and detailed patterns of cells within a specific cell type, providing a quantitative framework to better understand cell heterogeneity.

**Fig. 6:**
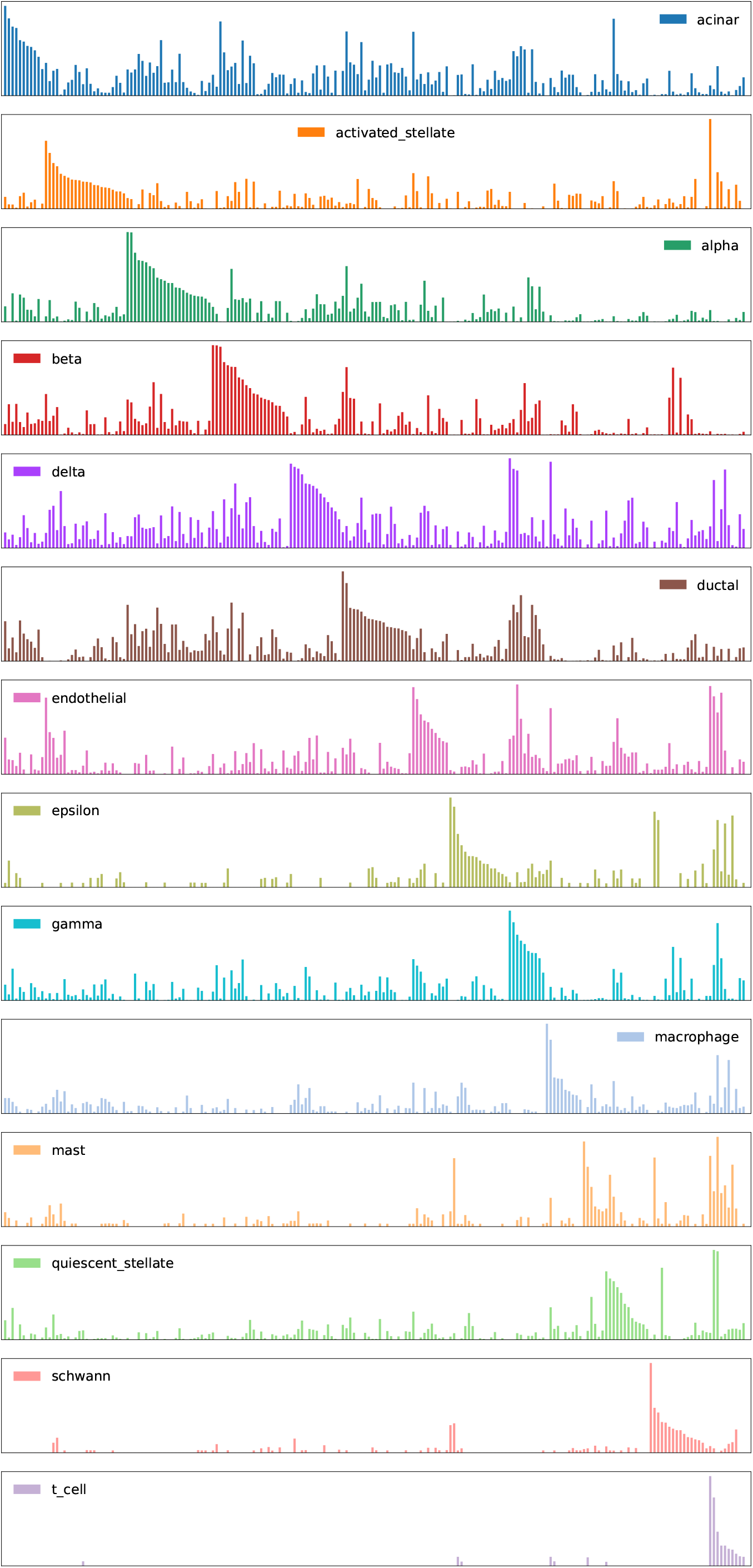
The cell type feature spectrum. The feature spectrum of cell types in the pancreas dataset, derived from the TF-IDF transformation of code frequency utilization, revealed distinct characteristics for each cell type.

## 4. Discussion

Large foundation models (FMs), pre-trained on extensive single-cell datasets, have recently achieved significant progress in the field of single-cell analysis (Yang et al. [2022], Cui et al. [2024a], Szalata et al. [2024]). Prominent examples of these cutting-edge models include scGPT (Cui et al. [2024a]) and scBERT (Yang et al. [2022]), which are widely utilized and studied in single-cell research and comparative analyses. The pre-training of these FMs, followed by fine-tuning on specific downstream tasks, enables the use of much smaller, application-specific datasets compared to training models from scratch. This sample efficiency is particularly advantageous in the biological domain, where clinical data or experimentally labeled datasets are often challenging and expensive to obtain, resulting in limited datasets for downstream applications.

Despite the significant advancements of foundation models (FMs) in single-cell analysis, several limitations and criticisms remain. For instance, Boiarsky et al. demonstrated that a simple regression model can outperform FMs in fine-tuning tasks for cell-type annotation, depending on the specific dataset (Boiarsky et al. [2023]). Similarly, Kedzierska et al. showed that FMs can underperform in clustering tasks compared to simpler neural network methods (Kedzierska et al. [2023]). These findings are consistent with our results, where supervised neural network approaches often achieved competitive or even superior performance compared to FMs.

A key factor contributing to these observations is the potential overfitting or inefficiency in fine-tuning FMs when the data distributions of pre-training and downstream tasks are not closely aligned (Varis and Bojar [2021]). FMs typically have a large number of trainable parameters. For example in our study, the total number of learnable parameters is approximately 6.5 million for scBERT and 51 million for scGPT. In comparison, SCLSC and scVQC have 1 million and 3 million parameters, respectively. Overfitting occurs when these models become excessively complex and large, causing them to memorize the training data rather than learning meaningful patterns. This results in poor generalization performance on unseen data. Supervised methods, specifically designed for labeled datasets, excel at capturing dataset-specific features and relationships, offering a tailored approach to the analysis.

Another challenge for FMs is their reliance on large-scale, high-quality pre-training datasets. Any biases or noise in these datasets can propagate through the models, reducing their generalizability to niche or highly specific single-cell datasets (Wang and Russakovsky [2023]). In contrast, supervised methods are task- and tissue-specific, enabling fine-tuned optimization that minimizes the impact of pre-training biases. This targeted learning approach allows supervised models to more effectively capture biologically meaningful structures and distinctions between cell types.

On the other hand, while FMs are designed to generalize across diverse datasets and tasks, this broad generalization can dilute their ability to fine-tune embeddings for specific datasets or tissues. Additionally, supervised models generally demand less computational effort compared to foundation models, which often involve a vast number of parameters. This reduced computational overhead not only makes supervised models competitive in terms of performance but also more accessible to researchers with limited computational resources. Our results further demonstrate that supervised approaches can outperform foundation models in terms of speed, sometimes by a significant margin.

In this study, we introduced scVQC, a cell type annotation framework that employs a split-Vector-Quantizer (split-VQ) to learn discrete embeddings of cells in a lower-dimensional space. Using codebooks generated from split-VQ, we can construct a novel representation of cells called the feature spectrum. The feature spectrum for each cell type is derived by applying the TF-IDF transformation to the utilization of codes within that cell type, analogous to identifying key words in documents. This method highlights features that are enriched or depleted for specific cell types, offering a powerful way to visualize overarching and detailed patterns within cell populations. Moreover, it provides a systematic approach to summarize complex single-cell RNA-seq data, potentially enabling the discovery of unique molecular signatures, rare cell types, and variations within heterogeneous populations, such as immune cells in PBMC datasets. Additionally, the feature spectrum serves as a bridge between computational analysis and biological interpretation, supporting hypothesis generation and deepening biological insights. Previous studies have applied the feature spectrum to characterize protein subcellular localization (Kobayashi et al. [2022]) and the epigenomic profiles of single cells (Cui et al. [2024b]). However, to the best of the authors’ knowledge, this study is the first to utilize the feature spectrum in the analysis of single-cell transcriptomic datasets.

## 5. Conclusion

In this study, we compared the effectiveness of foundation models (FMs) and supervised neural network approaches for the task of cell type prediction. Our findings showed that supervised approaches achieved performance comparable to FMs while requiring significantly less time and computational resources. Additionally, we introduced scVQC, a classifier based on split-vector quantization, which effectively learns cell representations for classification tasks. Moreover, scVQC can generate the cell type feature spectrum, a novel characterization of cell types derived from TF-IDF transformations of cell gene profiles quantized using the split-VQ method.

## 7. Code availability

The source code for scVQC is accessible at https://github.com/yusri-dh/scVQC

## 8. Data availability

We used 4 human single cell datasets in our study which can be accessed as follows:

- PBMC 68k (Zheng et al. [2017]): https://www.10xgenomics.com/resources/datasets/fresh-68-k-pbm-cs-donor-a-1-standard-1-1
- pancreas (Luecken et al. [2021]): https://figshare.com/ndownloader/files/22891151
- lung (Luecken et al. [2021]): https://figshare.com/ndownloader/files/24539942
- thymus (Park et al. [2020]): https://zenodo.org/record/5500511

All datasets were accessed and downloaded in June 2024.

## 9. Author contributions statement

Y.D.H. was responsible for the the data curation, analyses, and visualization, and writing the original draft of the manuscript.

Y.Z. contributed to study conceptualization, supervision, and manuscript editing. S.I. managed funding acquisition, project administration, supervision, and manuscript editing. All authors reviewed the manuscript.

## 11. Acknowledgments

The authors thank the anonymous reviewers for their valuable suggestions.

## References

[1] R. Boiarsky, N. Singh, A. Buendia, G. Getz, and D. Sontag. A deep dive into single-cell rna sequencing foundation models, Oct. 2023. URL http://dx.doi.org/10.1101/2023.10.19.563100.

[2] Z. A. Clarke, T. S. Andrews, J. Atif, D. Pouyabahar, B. T. Innes, S. A. MacParland, and G. D. Bader. Tutorial: guidelines for annotating single-cell transcriptomic maps using automated and manual methods. Nature Protocols, 16(6):2749–2764, May 2021. ISSN 1750-2799. doi: 10.1038/s41596-021-00534-0. URL http://dx.doi.org/10.1038/s41596-021-00534-0.

[3] H. Cui, C. Wang, H. Maan, K. Pang, F. Luo, N. Duan, and B. Wang. scgpt: toward building a foundation model for single-cell multi-omics using generative ai. Nature Methods, 21(8):1470–1480, Feb. 2024a. ISSN 1548-7105. doi: 10.1038/s41592-024-02201-0. URL http://dx.doi.org/10.1038/s41592-024-02201-0.

[4] X. Cui, X. Chen, Z. Li, Z. Gao, S. Chen, and R. Jiang. Discrete latent embedding of single-cell chromatin accessibility sequencing data for uncovering cell heterogeneity. Nature Computational Science, 4(5):346–359, May 2024b. ISSN 2662-8457. doi: 10.1038/s43588-024-00625-4. URL http://dx.doi.org/10.1038/s43588-024-00625-4.

[5] B. Dai, Z. Wang, and D. Wipf. The usual suspects? reassessing blame for vae posterior collapse, 2019. URL https://arxiv.org/abs/1912.10702.

[6] T. Dao, D. Y. Fu, S. Ermon, A. Rudra, and C. Ré. Flashattention: Fast and memory-efficient exact attention with io-awareness, 2022. URL https://arxiv.org/abs/2205.14135.

[7] J. Devlin, M.-W. Chang, K. Lee, and K. Toutanova. Bert: Pre-training of deep bidirectional transformers for language understanding, 2018. URL https://arxiv.org/abs/1810.04805.

[8] J. Ding and A. Regev. Deep generative model embedding of single-cell rna-seq profiles on hyperspheres and hyperbolic spaces. Nature Communications, 12(1), May 2021. ISSN 2041-1723. doi: 10.1038/s41467-021-22851-4. URL http://dx.doi.org/10.1038/s41467-021-22851-4.

[9] C. Eick, N. Zeidat, and Z. Zhao. Supervised clustering - algorithms and benefits. In 16th IEEE International Conference on Tools with Artificial Intelligence, TAI-04, page 774–776. IEEE Comput. Soc, 2004. doi: 10.1109/ictai.2004.111. URL http://dx.doi.org/10.1109/ICTAI.2004.111.

[10] D. P. Gomari, A. Schweickart, L. Cerchietti, E. Paietta, H. Fernandez, H. Al-Amin, K. Suhre, and J. Krumsiek. Variational autoencoders learn transferrable representations of metabolomics data. Communications Biology, 5(1), June 2022. ISSN 2399-3642. doi: 10.1038/s42003-022-03579-3. URL http://dx.doi.org/10.1038/s42003-022-03579-3.

[11] Y. D. Heryanto, Y.-z. Zhang, and S. Imoto. Predicting cell types with supervised contrastive learning on cells and their types. Scientific Reports, 14(1), Jan. 2024. ISSN 2045-2322. doi: 10.1038/s41598-023-50185-2. URL http://dx.doi.org/10.1038/s41598-023-50185-2.

[12] K. Z. Kedzierska, L. Crawford, A. P. Amini, and A. X. Lu. Assessing the limits of zero-shot foundation models in single-cell biology, Oct. 2023. URL http://dx.doi.org/10.1101/2023.10.16.561085.

[13] H. Kobayashi, K. C. Cheveralls, M. D. Leonetti, and L. A. Royer. Self-supervised deep learning encodes high-resolution features of protein subcellular localization. Nature Methods, 19(8):995–1003, July 2022. ISSN 1548-7105. doi: 10.1038/s41592-022-01541-z. URL http://dx.doi.org/10.1038/s41592-022-01541-z.

[14] M. D. Luecken, M. Büttner, K. Chaichoompu, A. Danese, M. Interlandi, M. F. Mueller, D. C. Strobl, L. Zappia, M. Dugas, M. Colomé-Tatché, and F. J. Theis. Benchmarking atlas-level data integration in single-cell genomics. Nature Methods, 19(1):41–50, Dec. 2021. ISSN 1548-7105. doi: 10.1038/s41592-021-01336-8. URL http://dx.doi.org/10.1038/s41592-021-01336-8.

[15] H. Naveed, A. U. Khan, S. Qiu, M. Saqib, S. Anwar, M. Usman, N. Akhtar, N. Barnes, and A. Mian. A comprehensive overview of large language models, 2023. URL https://arxiv.org/abs/2307.06435.

[16] A. v. d. Oord, O. Vinyals, and K. Kavukcuoglu. Neural discrete representation learning, 2017. URL https://arxiv.org/abs/1711.00937.

[17] J.-E. Park, R. A. Botting, C. Domínguez Conde, D.-M. Popescu, M. Lavaert, D. J. Kunz, I. Goh, E. Stephenson, R. Ragazzini, E. Tuck, A. Wilbrey-Clark, K. Roberts, V. R. Kedlian, J. R. Ferdinand, X. He, S. Webb, D. Maunder, N. Vandamme, K. T. Mahbubani, K. Polanski, L. Mamanova, L. Bolt, D. Crossland, F. de Rita, A. Fuller, A. Filby, G. Reynolds, D. Dixon, K. Saeb-Parsy, S. Lisgo, D. Henderson, R. Vento-Tormo, O. A. Bayraktar, R. A. Barker, K. B. Meyer, Y. Saeys, P. Bonfanti, S. Behjati, M. R. Clatworthy, T. Taghon, M. Haniffa, and S. A. Teichmann. A cell atlas of human thymic development defines t cell repertoire formation. Science, 367(6480), Feb. 2020. ISSN 1095-9203. doi: 10.1126/science.aay3224. URL http://dx.doi.org/10.1126/science.aay3224.

[18] Sza-lata, K. Hrovatin, S. Becker, A. Tejada-Lapuerta, H. Cui, Wang, and F. J. Theis. Transformers in single-cell omics: a review and new perspectives. Nature Methods, 21(8):1430–1443, Aug. 2024. ISSN 1548-7105. doi: 10.1038/s41592-024-02353-z. URL http://dx.doi.org/10.1038/s41592-024-02353-z.

[19] D. Varis and O. Bojar. Sequence length is a domain: Length-based overfitting in transformer models. In Proceedings of the 2021 Conference on Empirical Methods in Natural Language Processing. Association for Computational Linguistics, 2021. doi: 10.18653/v1/2021.emnlp-main.650. URL http://dx.doi.org/10.18653/v1/2021.emnlp-main.650.

[20] A. Wang and O. Russakovsky. Overwriting pretrained bias with finetuning data. In Proceedings of the IEEE/CVF International Conference on Computer Vision (ICCV), pages 3957–3968, October 2023.

[21] F. Yang, W. Wang, F. Wang, Y. Fang, D. Tang, J. Huang, H. Lu, and J. Yao. scbert as a large-scale pretrained deep language model for cell type annotation of single-cell rna-seq data. Nature Machine Intelligence, 4(10):852–866, Sept. 2022. ISSN 2522-5839. doi: 10.1038/s42256-022-00534-z. URL http://dx.doi.org/10.1038/s42256-022-00534-z.

[22] H. Zeng. What is a cell type and how to define it? Cell, 185 (15):2739–2755, July 2022. ISSN 0092-8674. doi: 10.1016/j.cell.2022.06.031. URL http://dx.doi.org/10.1016/j.cell.2022.06.031.

[23] G. X. Y. Zheng, J. M. Terry, P. Belgrader, P. Ryvkin, Z. W. Bent, R. Wilson, S. B. Ziraldo, T. D. Wheeler, G. P. McDermott, J. Zhu, M. T. Gregory, J. Shuga, L. Montesclaros, J. G. Underwood, D. A. Masquelier, S. Y. Nishimura, M. Schnall-Levin, P. W. Wyatt, C. M. Hindson, R. Bharadwaj, A. Wong, K. D. Ness, L. W. Beppu, H. J. Deeg, C. McFarland, K. R. Loeb, W. J. Valente, N. G. Ericson, E. A. Stevens, J. P. Radich, T. S. Mikkelsen, B. J. Hindson, and J. H. Bielas. Massively parallel digital transcriptional profiling of single cells. Nature Communications, 8(1), Jan. 2017. ISSN 2041-1723. doi: 10.1038/ncomms14049. URL http://dx.doi.org/10.1038/ncomms14049.

